# Tracing the ’Panda of the Sea’: Species-Specific qPCR Assays for eDNA Monitoring of Critically Endangered *Bahaba taipingensis* in the Pearl River Estuary

**DOI:** 10.64898/2026.07.20.739478

**Authors:** Yuzhuo Liao, Brandon Chao-Feng Szeto, Haohang Zhao, Haimei Lin, Guanlin Chen, Guang-Jie Zhou, Jiezhang Mo, Alicia Lee Sian Tan, Junjie Wang, Jack Chi-Ho Ip

## Abstract

Chinese Bahaba (*Bahaba taipingensis*, Sciaenidae) is a Critically Endangered, Grade I state-protected marine fish in China. Reliable, non-invasive monitoring tools are urgently needed to inform conservation and restocking efforts. We developed and validated three TaqMan probe–based qPCR assays targeting mitochondrial 12S rRNA, ND5, and control region (D-loop) loci for species-specific detection of *B. taipingensis*. Primer–probe sets were designed from complete mitochondrial genomes and evaluated *in silico* against GenBank and *in vitro* against tissue DNA from *B. taipingensis* and closely related sciaenids, plus positive eDNA samples; sensitivity was quantified using serial dilutions of synthetic target DNA. All three assays showed high specificity and consistent amplification; 12S rRNA assay (Bhb_12S) was selected for field screening based on superior low-copy sensitivity and minimal by-products. We applied the Bhb_12S assay to 414 seawater samples collected from 23 sites in western Hong Kong during 2025 surveys, detecting *B. taipingensis* eDNA in the September samples (0.71–2.85 copies per μL; mean 2.05), with positive detections confirmed by Sanger sequencing. Overall, these validated qPCR assays provide a robust molecular toolkit for non-invasive monitoring of *B. taipingensis* and will aid conservation planning, restocking evaluation, and long-term biodiversity assessments in the Pearl River Estuary and adjacent coastal waters.

## 1. Introduction

Chinese Bahaba (*Bahaba taipingensis*), a fish species within the family Sciaenidae, is endemic to the coastal waters of the East and South China Seas and is widely known as the “panda of the sea” due to its extreme rarity and immense cultural-economic value (Hu et al., 2025). As one of the largest members of the family Sciaenidae, it is particularly well known for its swim bladder, which is highly valued in traditional medicine and often commands prices comparable to gold, thus subjected the species to intense fishing pressure (Sadovy and Cheung, 2003). In addition, *B. taipingensis* preferentially inhabits nearshore coast environments and exhibits estuarine-dependent spawning migrations (entering estuaries to form large aggregations for reproduction), rendering it especially vulnerable to a combination of overexploitation, estuarine habitat degradation, pollution, coastal development, and reclamation—pressures that are particularly acute in heavily modified systems like the Pearl River Estuary (PRE), which remains one of its last major spawning refuges (Sadovy and Cheung 2003; Chen et al. 2025). As a result of these combined pressures, wild populations of *B. taipingensis* have experienced a dramatic decline over recent decades, with systematic assessments revealing substantial range contraction (approximately 70% reduction in historical distribution), significant decreases in maximum body size and catching rate (Chen et al. 2025). Reports of captures of wild individuals have become exceedingly rare, underscoring the urgent need for effective conservation and monitoring.

A series of conservation and management measures have been implemented to protect this species. China has listed *B. taipingensis* as a second-class species on the List of State Key Protected Wild Animals in 1988, and upgraded it to first-class (Grade I) protection in 2021 — a significant elevation that places it in the same conservation category as the giant panda (Hu et al., 2025). Notably, *B. taipingensis* is currently the only exclusively marine fish species granted first-class national protection status in China. At the international level, the species was classified as Critically Endangered (CR) by the International Union for Conservation of Nature (IUCN) in 2006 (Wang et al., 2009), further highlighting its global conservation significance. Substantial efforts have also been made from ecological perspective. In 2005, a nature reserve dedicated to *B. taipingensis* was established in Dongguan, Guangdong Province; more recently, the first successful artificial reproduction of the species was achieved in 2021, paving the way for intensive captive breeding and stock enhancement programs (Hu et al., 2025). These programs now release thousands of juveniles annually into the PRE system (e.g., 500 bred in Shenzhen Bay, Shenzhen in 2023; 1,320 bred in Nansha, Guangzhou in 2025), providing renewed hope for population recovery but also highlighting the urgent need for reliable post-release monitoring.

Despite these protections and advances in captive breeding, progress in understanding contemporary wild populations has remained limited. Moreover, with increasing stock enhancement efforts, there is a growing need for non-invasive ecological studies— particularly population monitoring—to evaluate restocking success, track dispersal and habitat use, and provide feedback for adaptive management. The Pearl River Estuary represents a historically important region for *B. taipingensis* (Sadovy and Cheung 2003) and is therefore a priority area for population assessment. Such studies can provide critical guidance for breeding and release programs, supporting the evaluation of ongoing conservation actions (such as restocking) and informing future strategies, including potential expansion of protected areas or targeted habitat restoration.

As with many marine fish surveys, monitoring of *B. taipingensis* has traditionally relied on conventional fisheries-based approaches, such as trawling (Sadovy and Cheung 2003). However, these methods are often logistically demanding, financially costly, and destructive to benthic habitats and non-target organisms, and—crucially—ineffective for detecting rare, low-density, or spatially patchy populations like *B. taipingensis*. In the Pearl River Estuary (PRE), their limitations are exacerbated by complex hydrological conditions (e.g., high sediment loads, variable salinity, and strong tidal mixing), which further reduce encounter probabilities in capture-based surveys (Zou et al., 2020; Ip et al., 2024). Under these circumstances, there is a clear and urgent need for an effective, non-invasive, and sensitive monitoring tools to support conservation-oriented assessments of this critically endangered species.

Environmental DNA (eDNA) analysis has emerged as a powerful, non-destructive alternative to traditional surveys, enabling detection of species presence through genetic material (e.g., mucus, feces, gametes, or sloughed cells) shed into the environment (Taberlet et al., 2012; Thomsen and Willerslev, 2015). To date, eDNA-based approaches have been widely validated as effective tools for biodiversity surveys across a range of ecosystems, including terrestrial (Leempoel et al., 2020), freshwater (Nakagawa et al., 2018; He et al., 2022), and marine environments (Thomsen et al., 2012; Stat et al., 2019). The integration of eDNA with real-time quantitative PCR (qPCR) has further expanded its utility, particularly for targeted detection of rare or elusive taxa (Klymus et al., 2015). By employing species-specific primer-probe sets and highly sensitive chemistries, qPCR can reliably amplify trace DNA amounts (often at picogram levels or lower), outperforming conventional PCR or metabarcoding in scenarios with low target abundance or high non-target competition (LeBlanc et al., 2020). Metabarcoding with universal primers, while excellent for community profiling, is prone to amplification bias and may miss extremely rare species due to primer competition or PCR inhibition (Pawluczyk et al., 2015); thus, the high sensitivity and specificity make qPCR-based eDNA approaches particularly suitable for the detection of such species (Xia et al., 2021). In contrast, qPCR-based assays have proven highly effective for low-abundance endangered fish, such as the extremely rare bitterling fish (Otsuki et al., 2023) and white sturgeon (Crossman et al., 2024).

Despite these advances, species-specific qPCR assays remain unavailable for the Chinese Bahaba (*B. taipingensis*), hindering the implementation of accurate, non-invasive eDNA surveys for this ecologically and culturally significant species. In this study, we aimed to (1) develop and rigorously validate species-specific TaqMan qPCR assays targeting mitochondrial loci for sensitive detection of *B. taipingensis* eDNA, and (2) apply the most robust assay to field surveys in western Hong Kong waters (part of the PRE system) to assess species presence. These results establish a novel molecular toolkit that enables advanced, non-invasive monitoring and directly supports conservation efforts—including evaluation of restocking programs—for this critically endangered flagship species.

## 2. Materials and Methods

### 2.1 Primer development and *in silico* testing

Compared with nuclear DNA, mitochondrial DNA (mtDNA) is generally a more suitable molecular marker for eDNA assays due to its substantially higher cellular copy number (Rees et al., 2014). Based on previous eDNA studies involving fish taxa in the marine environment (Baker et al., 2018; Tsuji et al., 2018; Zhang et al., 2020), we selected three mtDNA regions — 12S-Ribosomal RNA (12S), NADH dehydrogenase 5 (ND5), and Control Region (D-loop) — for species-specific primer and probe development. Complete mitochondrial genome sequences of *B*. *taipingensis* were retrieved from NCBI GenBank. Additionally, based on studies about phylogenetic analysis of *B*. *taipingensis* (Zhao et al., 2015), mitochondrial genomes from ten phylogenetically related species in family Sciaenidae (*Collichthys lucidus*, *C*. *niveatus*, *Larimichthys crocea*, *L*. *polyactis*, *Parapristipoma trilineatum*, *Dendrophysa russelii*, *Pennahia argentata*, *Miichthys miiuy*, *Nibea albiflora*, and *N*. *coibor*), all of which potentially overlap with *B. taipingensis* in geographic distribution, were obtained for comparison. All sequence data used for primer development of *B*. *taipingensis* are listed in Table S1.

Multiple sequence alignments were conducted in Geneious Prime v2025.0.2 (Biomatters, New Zealand) to identify highly variable interspecific regions within each target locus. Primer and minor-groove binding (MGB) probe design, along with preliminary screening, were performed using Primer3.0 (Untergasser et al., 2012) under criteria specifying optimal melting temperature of 60 °C, GC content around 40-60 %, and minimal self-complementarity. The primer-probe sets for each target region were further *in silico* evaluated using NCBI Primer-BLAST to confirm amplification specificity and eliminate potential off-target matches (2 or less mismatches to the primer). Details of the three selected primer-probe sets are provided in Table 1.

**Table 1.**
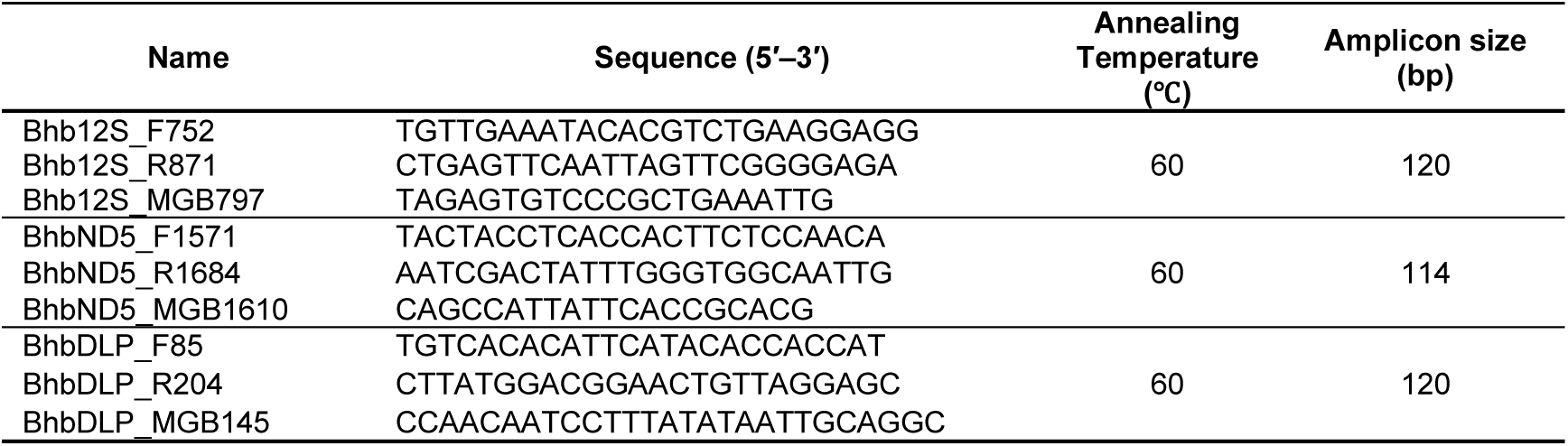
Details of the new primer-probe sets targeting the mitochondrial genome of Chinese Bahaba.

### 2.2 Specificity and sensitivity testing

We evaluated the specificity of the designed *B*. *taipingensis* primers by conducting PCR using *B. taipingensis* tissue DNA (fin clips) as positive control and six closely related sciaenid species: *L. crocea*, *C. lucidus*, *Johnius. belangerii*, *Pennahia. argentata*, *N. albiflora*, and *N*. *squamosa*. For *B. taipingensis*, DNA extraction and PCR analysis were conducted in co-author’s eDNA Lab at South China Normal University (Figure 1), using tissue collection from the Chinese Bahaba breeding farm in Huizhou, Guangdong province, China. Tissues of the other commercial species were obtained from Hong Kong’s fish market. Genomic DNA was extracted using the DNeasy Blood & Tissue Kit (Qiagen, Germany) following the manufacturer’s instructions, and subsequently, PCR reactions were performed in a total volume of 25 μL, consisting of 1 μL each of forward and reverse primers (10 µM), 12.5 μL of Taq PCR Master Mix (Qiagen, Germany), 9.5 μL of nuclease-free water, and 1 μL of DNA template. The thermal cycling program included an initial denaturation at 94 °C for 5 min, followed by 35 cycles of denaturation at 94 °C for 30 s, annealing at 57 °C for 45 s, and extension at 72 °C for 1 min, with a final extension at 72 °C for 8 min. PCR products were visualized on 2 % agarose gel electrophoresis to identify the successful and specific amplicon of *B. taipingensis*.

**Figure 1.**
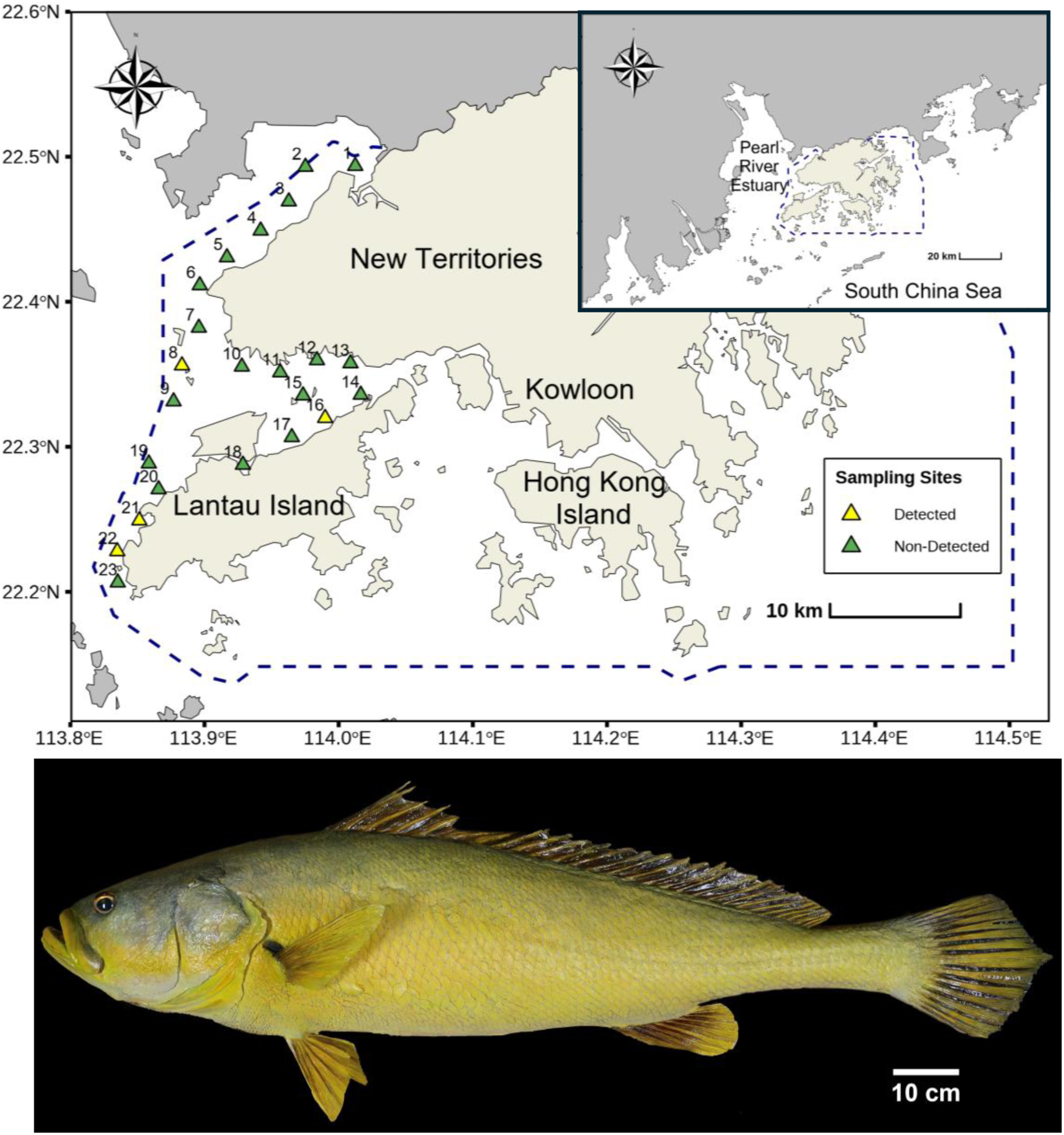
Map of western Hong Kong showing the 23 sampling sites used in this study (green, upper panel). Four sites yielded Chinese bahaba eDNA detections (yellow): Sha Chau and Lung Kwu Chau Marine Park (S8), The Brothers Marine Park (S16), Tai O (S21), and Southwest Lantau Marine Park (S22). Lower panel: photograph of an adult Chinese bahaba specimen (total length 152 cm) from the Bahaba Nature Reserve, Dongguan.

We also evaluated primer performance using positive eDNA samples collected from two sources: (1) an aquaculture tank containing *B. taipingensis* and (2) environmental samples from survey sites spiked with *B. taipingensis* tissue DNA through a ten-fold serial dilution (10⁻¹ to 10⁻⁴ ng/µL) within a natural eDNA background. Real-Time PCR (qPCR) was performed with on a StepOnePlus Real-Time PCR System (Applied Biosystems, USA). Reaction setup was conducted in a pre-PCR laboratory dedicated to preventing amplicon contamination. Each 20-µL reaction consisted of 10 µL TaqMan^™^ Environmental Master Mix 2.0 (Applied Biosystems, USA), 0.5 µL of each forward and reverse primer (10 µM), 0.5 µL of probe (10 µM), 6.5 µL nuclease-free water, and 2 µL of eDNA template. Thermal cycling conditions began with 50 °C for 2 min, 95 °C for 2 min, followed by 45 cycles of denaturation at 95 °C for 15 s and 60 °C for 1 min.

All positive PCR and qPCR products from above testing were purified and submitted to BGI Health (HK) Company Limited (Hong Kong, China) for Sanger sequencing. Resulting sequences were inspected in Geneious and compared against the NCBI GenBank database (Aug. 2025 version) using BLAST to confirm species identity, with *B. taipingensis* identified as the best match based on similarity > 98% and E-value threshold of 1e-30.

The limit of detection (LOD) and limit of quantification (LOQ) for each primer pair were determined from standard curves generated from qPCR assays (consistent setting with positive eDNA testing qPCR) of a serial dilution of synthetic target DNA (Taihe Biotechnology, China), ranging from 1 to 1×10^6^ copies per μL. LOD and LOQ weredetermined based on Ellison et al. (2006) and Chen et al. (2022), i.e., the LOD as the lowest concentration of the serial dilution producing over half positive replicate qPCR results, and the LOQ as the lowest concentration of the serial dilution at which all replicates are positive. ; the detection threshold (Ct) was defined for each primer set based on the highest Ct value observed among positive reactions in the LOD/LOQ tests.

### 2.3 Field survey and qPCR analysis

To investigate the migration of *B. taipingensis* across Hong Kong waters, we conducted six surveys during the migration season (March–June) and additional surveys before (January) and after (September) the migration period in 2025. Water samples were collected from 23 evenly distributed sites in the western Hong Kong waters, covering Mai Po Nature Reserve and three marine parks: Sha Chau and Lung Kwu Chau Marine Park, The Brothers Marine Park, and Southwest Lantau Marine Park (Figure 1). At each site, three 1-L surface seawater replicates and one 1-L negative control (sterile distilled water) were collected. Each seawater replicates were collected twice (500 mL per collection) using plastic water sampler, transferred into sterile sampling bags, and immediately stored on ice until filtration. All sampling equipment was sterilized before each survey by soaking in 1% bleach for 15 min, rinsing with sterile distilled water, and placing under UV light for 30 min; water sampler was sprayed with 1% bleach and cleaned with distilled water between sites as well during each survey. Following transportation to the laboratory, each sample was filtered through 47-mm mixed cellulose ester (MCE) filter with a pore size of 0.45 µm (Millipore, USA) inside a dedicated UV-sterilized eDNA hood within 6 hours. Filters were transferred to sterile Falcon petri dishes and immediately stored at -80 °C until DNA extraction.

eDNA was extracted using the DNeasy PowerSoil Pro Kit (Qiagen, Germany) following the manufacturer’s protocol in the laminar flow hood with UV sterilization (Ip et al., 2024); eDNA samples were processed in designated clean room spaces away from where the fish tissue was processed. DNA quantity was examined using Qubit 3.0 fluorometer (Thermo Fisher Scientific, USA). qPCR assays were performed followed by the method described in Section 2.2. Apart from site blank controls, each run included two blanks (nuclease-free water) as negative controls and two reactions containing *B. taipingensis* tissue DNA as positive controls. A site was considered positive if at least one site replicate showed a clear amplification curve and a Ct value below the predefined threshold. For positive reactions, only quantitative results calculated by a standard curve with R² > 0.98 and an amplification efficiency of 90–110% were accepted. Positive qPCR products were purified and submitted for Sanger sequencing, and the resulting sequences were identified using BLAST against the GenBank database.

## 3. Results

### 3.1 Primer specificity and sensitivity

Three selected primer-probe sets (namely: Bhb_12S, Bhb_ND5 and Bhb_DLP) demonstrated high specificity to *B. taipingensis*. Both *in-silico* and *in-vitro* tests confirmed the specificity of primers, with no off-target amplification detected in Primer-BLAST and testing PCR results (Figure 2A-C). All three primers also exhibited high sensitivity, as confirmed by qPCR results from positive eDNA samples (Figure 2A-C). The LOD was determined to be 1 copy/μL for all three primer sets; while the LOQ was 10 copies/μL for three primer sets.

**Figure 2.**
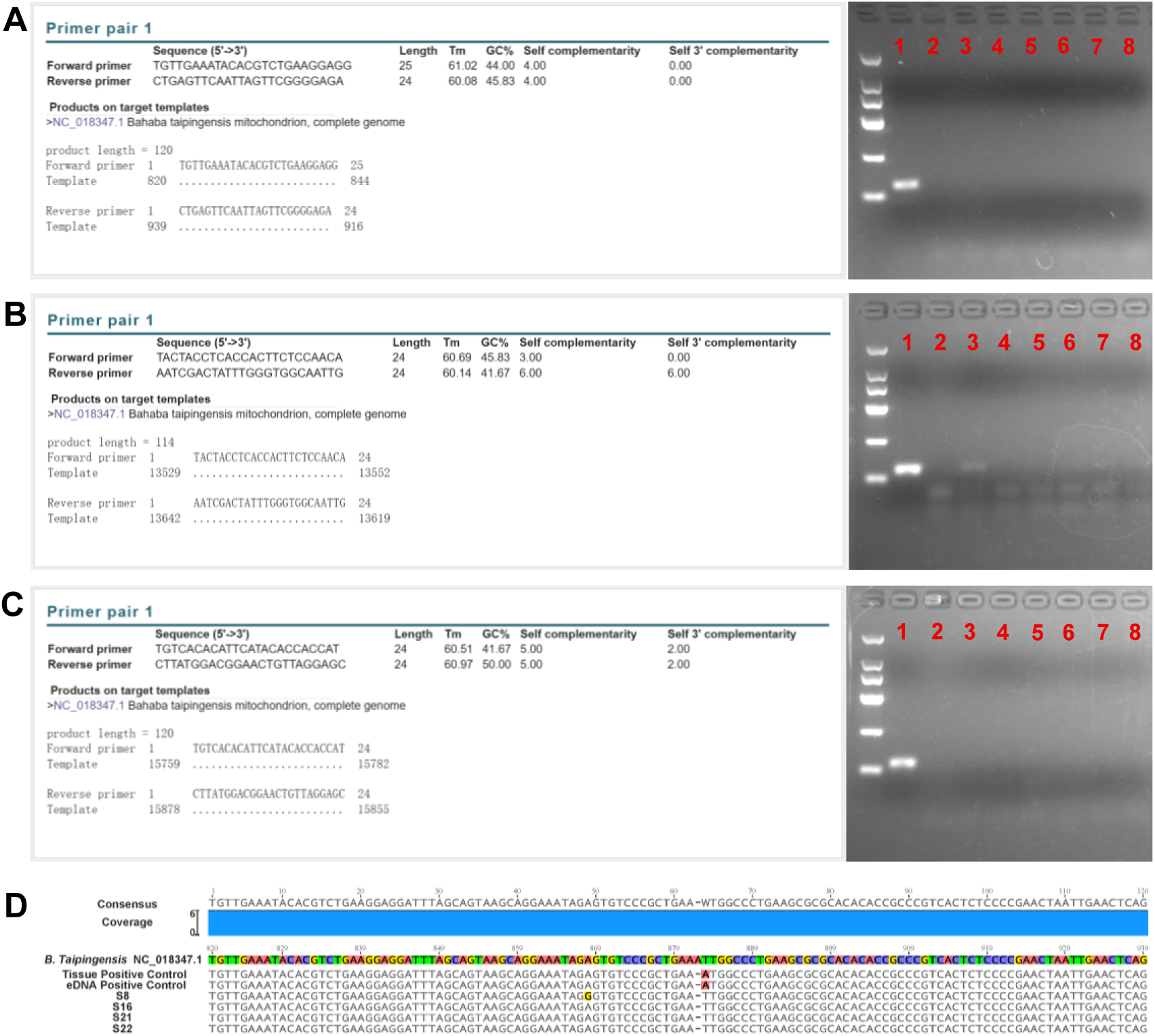
*In silico* and *in vitro* specificity testing of the three species-specific qPCR assays for *Bahaba taipingensis*. *In-silico* and *in-vitro* test result of three selected primers. (**A**) Bhb12S; (**B**) BhbND5; (**C**) BhbDLP. Template used for each *in-vitro* PCR test (from #1 to #8 in gel electrophoresis): *B. taipingensis* tissue, Nuclease-free water, *Larimichthys crocea* tissue, *Collichthys lucidus* tissue, *Johnius belangerii* tissue, *Pennahia argentata* tissue, *Nibea albiflora* tissue, and *N. squamosa* tissue. (**D**) Multiple alignment result of positive controls and positive samples (S8, S16, S21, S22 during Sep 2026 survey) with reference sequence of *B. taipingensis* from NCBI (Accession number: NC_018347.1).

Based on the testing qPCR results, the “detection threshold” (Ct) were set at 41.96 for Bhb_12S, 40.38 for Bhb_ND5, and 42.06 for Bhb_DLP, qPCR result of field sample with Ct value higher than the “detection threshold” will be discard as false positive; which ensuring reliable detection of *B. taipingensis* eDNA. Although all primer pairs showed good performance, Bhb_12S was selected for field screening due to its superior performance in LOD/LOQ tests for trace eDNA (<100 copies/μL) and a lower likelihood of generating by-products (Figure 3, Table S2).

**Figure 3.**
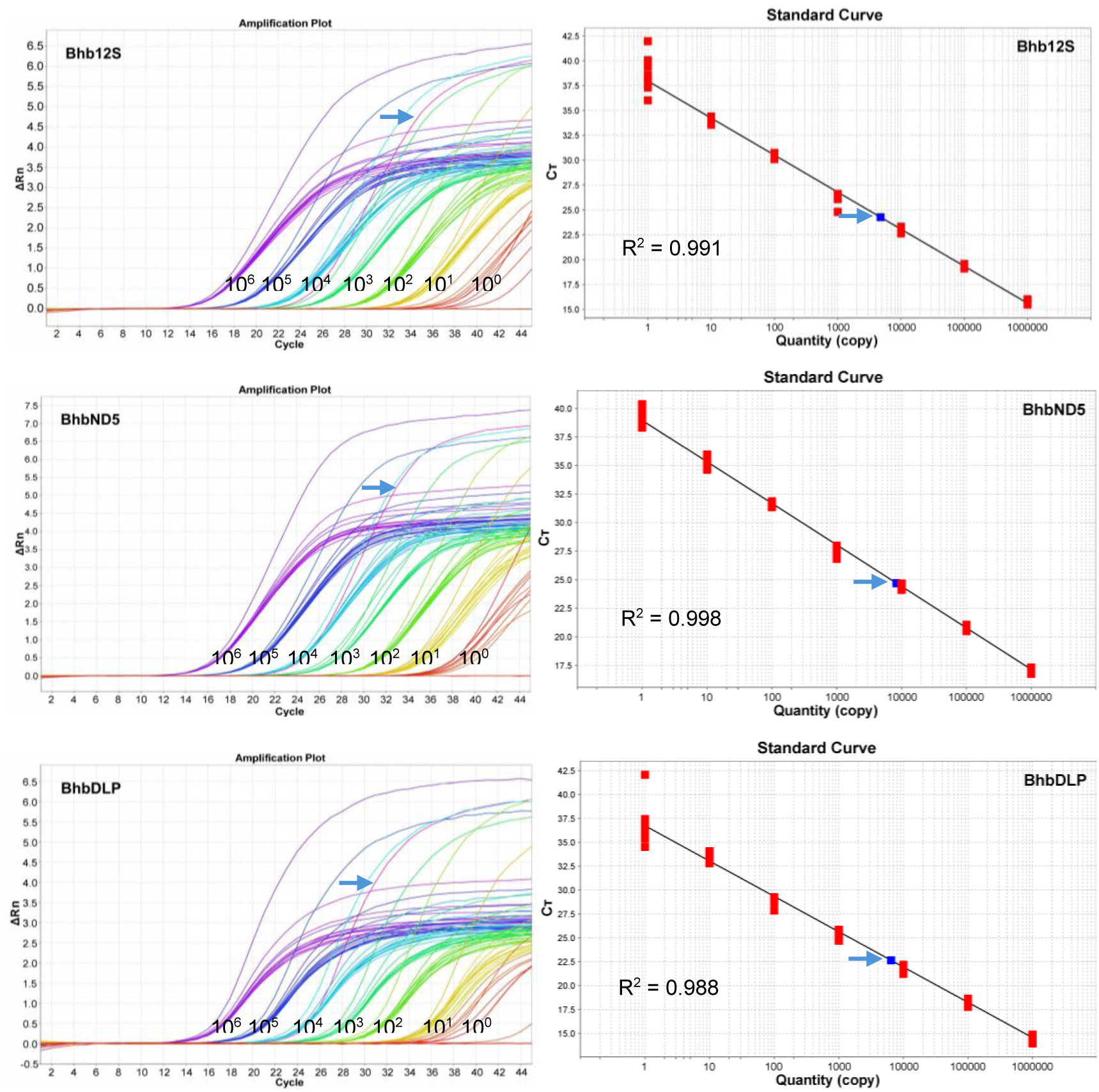
Limit of detection & limit of quantification test results of Three developed primer pairs. Left: amplification plot of the synthetic DNA serial dilution, with concentration ranging from 10^6^ to 10^0^ copies per μL; right: standard curve of each test, with the generated linear regression; blue arrows indicate result of qPCR positive control.

### 3.2 Detection of Chinese Bahaba eDNA in study area

To investigate whether *B. taipingensis* migrates across Hong Kong waters, we collected 414 seawater samples from 23 sites across western Hong Kong waters in 2025. During the migration period in 2025, *B. taipingensis* eDNA was not detected; however, interestingly, positive signals were identified at four out of the 23 sites (S8, S16, S21, and S22; Figure 2D) during the September 2025 survey. The concentration of *B. taipingensis* eDNA in positive reactions ranged from 0.71 to 2.85 copies per μL, with an average of 2.05 copies per μL(Table S3), with no amplification in the corresponding site blank and qPCR negative controls. Sanger sequencing of the positive samples confirmed the species identity of the qPCR products, with all sequences matching *B. taipingensis*. The detailed alignment results of positive samples with *B. taipingensis* are shown in Figure 2D.

## 4. Discussion

The development of species-specific primers for eDNA detection represents a critical advancement in monitoring critically endangered species like the Chinese Bahaba *B. taipingensis*, particularly in regions where traditional survey methods are ineffective due to the species’ rarity and cryptic nature. In this study, we present the first validated qPCR assays targeting mitochondrial regions (12S, ND5, and D-loop) for *B. taipingensis* eDNA, filling a key gap in conservation tools for this IUCN Critically Endangered fish.

### 4.1 Primer development and validation

Developing species-specific primers requires careful selection of target regions and reference sequences from phylogenetically and ecologically relevant species to ensure specificity and minimize off-target amplification (Tsuji et al. 2018; Hong et al. 2025; Osathanunkul and Suwannapoom 2025). Here, we aligned mitochondrial genomes from *B. taipingensis* and ten co-occurring Sciaenidae species, focusing on variable interspecific sites within the 12S, ND5, and D-loop loci. This approach, combined with *in silico* evaluation using NCBI Primer-BLAST, identified primer-probe sets with no significant matches to non-target taxa under stringent criteria (≤2 mismatches).

Rigorous validation of species-specific primers is essential for their application in eDNA surveys. In this study, three primer–probe sets were evaluated through a combination of *in-silico* screening and *in-vitro* testing using DNA from multiple sources. *In-silico* assessments against online databases confirmed the absence of non-target amplification under stringent matching criteria; laboratory experiments using both tissue-derived DNA and eDNA samples from *B. taipingensis* demonstrated reliable amplification performance, and the use of positive eDNA samples enabled validation under conditions that more closely reflect field surveys. In vitro specificity tests against six closely related sciaenids (including the sister species *L. crocea*, *C. lucidus*) showed no cross-amplification, while sensitivity assays on serially diluted synthetic DNA yielded low LOD and LOQ values. The comparable analytical sensitivities observed among the three assays provided essential information for selecting the most appropriate primer set for subsequent field applications and establish a robust methodological foundation for downstream eDNA surveys. Our validation aligns with level 4 of the five-level framework by Thalinger et al. (2021), incorporating *in silico*, *in vitro*, and preliminary field testing; achieving level 5 would require broader field-based detection probability assessments.

### 4.2 Field application and ecological insights

These validated assays enabled the first successful detection of *B. taipingensis* eDNA in natural seawater samples, extending the laboratory findings into real-world application. Using the Bhb_12S assay, we identified positive signals at four sites (S8: Sunny Bay, S16: Tsz Kan Chau, S21: Tai O, S22: Tsing Lam Kok) in western Hong Kong waters during September 2025—post-migration season and shortly after a documented restocking event of approximately 1,300 individuals into the Pearl River Estuary on 8 August 2025(from online report, http://cxzg.chinareports.org.cn/cxzg/news/66256.html). Notably, Tai O site (S21), a historically important fishing ground for the species (Sadovy and Cheung 2003), exhibited the highest detection frequency and lowest Ct values (high copy number of eDNA). All positive samples had low eDNA concentrations (0.71–2.85 copies per μL, average 2.05), with some of them below the LOD for Bhb_12S (1 copies/μL) but all Ct values of them were below the threshold (41.96). Such low-level signals are common in eDNA monitoring of rare species and should not be dismissed, as they may indicate genuine presence (Kralik and Ricchi, 2017; Klymus et al., 2020).

Given *B. taipingensis*’ extreme rarity—driven by overfishing and habitat loss—these detections provide novel evidence of potential habitat use in Hong Kong waters, informing conservation strategies like marine protected areas and restocking evaluations (Robinson et al. 2019; Hong et al. 2025). Consistent with previous studies, species-specific qPCR-based eDNA monitoring provides a sensitive and informative tool for evaluating the outcomes of restocking or eradication projects (Osathanunkul and Suwannapoom 2023). Overall, the absence of detections during the peak migration period (March–June), along with no amplification in negative controls and Sanger sequencing confirmation of amplicon identity, strongly supports the authenticity of these signals and rules out contamination. Additionally, with no detections in the PRE region in 2025 (unpublished data), these results suggest that the early warning signal associated with the decline in adult *B. taipingensis* may not be replicated every year, highlighting the urgent need for conservation efforts.

### 4.3 Significance of eDNA for monitoring endangered species

Currently, the probability of capturing *B. taipingensis* using conventional fisheries-dependent or observation-based methods is extremely low, rendering traditional surveys inefficient or impractical. In contrast, eDNA monitoring provides a highly sensitive and non-invasive alternative that allows the detection of species presence without physical capture or disturbance to individuals. As demonstrated in our study, meaningful distribution information can be obtained with relatively limited sampling effort, enabling spatially extensive surveys within a short time frame. These characteristics make the eDNA method particularly suited for the management and conservation of endangered species, where minimizing anthropogenic impacts while maximizing detection probability is a primary concern.

Moreover, eDNA-based surveys facilitate repeated sampling and long-term monitoring, offering a practical framework for assessing population persistence, responses to restocking projects, or changes in distribution over time. Similar eDNA approaches have proven effective for other endangered fish, such as cetacean in PRE and sturgeon in the Yangtze, highlighting its utility in biodiversity hotspots (Yu et al., 2021; Ushio et al., 2025).

### 4.4 Limitations and future directions

While qPCR eDNA assays excel in sensitivity, limitations include challenges in quantifying abundance due to factors like DNA shedding variability, degradation, and transport (Harrison et al., 2019; Sahu et al., 2025). Our low-copy detections underscore the risk of false negatives at trace levels, emphasizing the need for multi-replicate sampling and integration with traditional data (Ficetola et al., 2016). The mismatch between the restocking time in August and the detection of eDNA in September suggests that *B. taipingensis* may have entered Hong Kong’s western waters, including Sunny Bay (S8 site), shortly after release. To achieve full level 5 operational validation (Thalinger et al., 2021), future studies should quantify detection probabilities in known occupied and unoccupied sites.

Despite these limitations, this study presents the first species-specific qPCR assay targeting *B. taipingensis* eDNA, filling a critical methodological gap for monitoring and conserving this extremely rare species. The successful detection of *B. taipingensis* eDNA in field samples demonstrates the practical applicability of the assay in real environmental conditions.

When integrated with other species-specific or metabarcoding eDNA assays, the assay developed here could be incorporated into broader monitoring frameworks, enabling species-level detection of multiple threatened or ecologically important taxa. Such integration has the potential to enhance the efficiency and accuracy of biodiversity monitoring programs in data-limited coastal systems like the Pearl River Estuary (Zou et al., 2020; Ip et al., 2024; Chen et al., 2025), improving our understanding of habitat and species co-occurrence for better management practices in the future (e.g., Chinese Bahaba conservation regions and restocking locations).

## Supplementary materials

Supplementary data to this article can be found online at XXXX.

## Author Contributions

Y.L. developed and validated the assay, analyzed the data, and drafted the manuscript. B.C.F.S., H.Z., H.L., and G.C. performed fieldwork, processed eDNA samples, and curated the data. G.J.Z., J.M., and A.L.S.T. provided resources and critically reviewed the manuscript. J.W. and J.C.H.I. supervised the project, secured funding, and revised the manuscript. All authors approved the final version.

## Acknowledgements

The authors would like to acknowledge the person who have facilitated eDNA sample collection and molecular experiment for this project, specifically from the following research assistants and graduate students: Sandy Sze Man Yau, Phoebe Pui Ching Leung, Jujing Wang, Guiyu Tan, Wenjun Chen, Jinsheng Xiao.

## Data availability statement

Sanger sequence data for tissue and eDNA samples have been deposited in Figshare with the accession number: DOI/10.6084/m9.figshare.31558042.

